# nf-core/taxprofiler: highly parallelised and flexible pipeline for metagenomic taxonomic classification and profiling

**DOI:** 10.1101/2023.10.20.563221

**Authors:** Sofia Stamouli, Moritz E. Beber, Tanja Normark, Thomas A. Christensen, Lili Andersson-Li, Maxime Borry, Mahwash Jamy, nf-core community, James A. Fellows Yates

## Abstract

Metagenomic classification tackles the problem of characterising the taxonomic source of all DNA sequencing reads in a sample. A common approach to address the differences and biases between the many different taxonomic classification tools is to run metagenomic data through multiple classification tools and databases. This, however, is a very time-consuming task when performed manually - particularly when combined with the appropriate preprocessing of sequencing reads before the classification.

Here we present nf-core/taxprofiler, a highly parallelised read-processing and taxonomic classification pipeline. It is designed for the automated and simultaneous classification and/or profiling of both short- and long-read metagenomic sequencing libraries against a 11 taxonomic classifiers and profilers as well as databases within a single pipeline run. Implemented in Nextflow and as part of the nf-core initiative, the pipeline benefits from high levels of scalability and portability, accommodating from small to extremely large projects on a wide range of computing infrastructure. It has been developed following best-practise software development practises and community support to ensure longevity and adaptability of the pipeline, to help keep it up to date with the field of metagenomics.

## 2 Introduction

Whole-genome, metagenomic sequencing offers strong benefits to the taxonomic classification of DNA samples over targeted approaches (Eloe-Fadrosh et al. 2016; Florian P. Breitwieser, Lu, and Salzberg 2019). While metabarcoding approaches targeting the 16S rRNA or other marker genes are widely used due to low cost and large, diverse reference databases (Yilmaz et al. 2014; Lynch and Neufeld 2015), metagenomic approaches have been gaining popularity with the increasingly lower costs of, for example, shotgun sequencing. These metagenomic analyses with whole microbial genome as references have been shown to provide a similar level of taxonomic resolution (Hillmann et al. 2018). However they also have the added benefit of having greater reusability potential of the data, such as for whole genome and/or functional classification (Sharpton 2014; Quince et al. 2017).

Taxonomic classifiers (sometimes referred to as taxonomic binners) aim to identify the original ‘taxonomic source’ of a given DNA sequence (Ye et al. 2019; Meyer et al. 2022; Govender and Eyre 2022). In metagenomics, this typically consists of comparing millions of DNA reads (sequenced DNA molecules) against hundreds or thousands of reference genomes either via sequence alignment or ‘k-mer matching’ (Sharpton 2014; Sun et al. 2021). The reference genome with the most similar match to the read is then considered the most likely original ‘source’ organism of that sequence. In this article we will also refer to ‘taxonomic profilers’. We consider these as classifiers that also try to infer sequence abundance (i.e. re-assignment of counts to the most likely source based on the distribution of other hits) or biological relative abundance of the organism in the original sample (by coverage of expected marker genes, copy number estimations etc.), in addition to the simple read classification (Nayfach and Pollard 2016). We will use classifiers and profilers interchangeably throughout the publication.

Having to identify the original source of the many DNA sequences out of the many reference genomes in a time and computationally efficient manner is a difficult problem. In many cases biologists are not just interested as to which organism of each DNA sequence comes from, but also in using this information to infer the original ‘cellular’ or natural) abundance of each organism of the given environment - something that is very difficult due to the biases inherent to DNA extraction and sequencing. Therefore a plethora of tools have been developed to address these challenges, all with their own biases and specific contexts (Sczyrba et al. 2017; Meyer et al. 2022). Furthermore, each tool often produces tool-specific output formats making it difficult to efficiently cross compare results. Thus, no established ‘gold standard’ classifier tool or method currently exists.

One solution to addressing the problem of choice among the range of different tools is to run all of them in parallel, and cross compare the results. This can be useful both for benchmarking studies (e.g. Sczyrba et al. 2017; Meyer et al. 2022), but also to build consensus profiles whereby confidence of a particular taxonomic identification can be increased when it is detected by multiple tools (McIntyre et al. 2017; Ye et al. 2019).

A second challenge in taxonomic classification (and arguably a larger one) is a question of databases. As with tools, there is no one set ‘gold standard’ database. Different questions and contexts require different databases, such as when a researcher wants to search for both bacterial and viral species in samples, but as an extension of this, taxonomic classifiers often will need different settings for each database. Furthermore, as genomic sequencing becomes cheaper and more efficient, the number of publicly available reference genomes is rapidly increasing (Nasko et al. 2018). Consequently, the size of reference databases of taxonomic classifiers is also growing, often outpacing the computational capacity available to researchers. In fact, while this was one of the main motivations behind classifiers such as Kraken2 (Wood, Lu, and Langmead 2019), these algorithmic techniques are already becoming insufficient (Wright, Comeau, and Langille 2023).

Finally, with the decrease of costs, the possibility for larger and larger metagenomic sequencing datasets increases, leading to increasing sample sizes in studies. This is exemplified by the doubling of the number of metagenomes on the European Bioinformatic Institute’s MGnify database within just two years (Mitchell et al. 2019).

Altogether this highlights the need for methods to efficiently profile many samples using many tools. Manually setting up bioinformatic jobs for classification tasks for each database and settings against different tools on traditional academic computing infrastructure (e.g. high performance computing clusters or ‘HPC’ clusters) can be very tedious. Additionally, particularly for very large sample sets, there is increasing use of cloud platforms that have greater scalability than traditional HPCs. Being able to reliably and reproducibly execute taxonomic classification tasks across infrastructure with minimal intervention would therefore be a boon for the metagenomics field.

In recent years, workflow managers such as Nextflow (Di Tommaso et al. 2017) or Snakemake (Mölder et al. 2021) have become highly popular in bioinformatics. These frameworks provide for developers robust workflow execution with different HPC scheduling tools and software provisioning systems, ensuring maximum portability and efficient in different computational contexts. While a range of metagenomic pipelines already exist (a non-exhaustive list being for example, StaG-mwc by Boulund et al. 2023; MetaMeta by Piro, Matschkowski, and Renard 2017; TAMA by Sim et al. 2020; UGENE by Rose et al. 2019; and Sunbeam by Clarke et al. 2019), few leverage workflow managers to make multi-step workflows easier to use in HPC or cloud infrastructure. Furthermore, often these pipelines aim to carry out multiple different types of metagenomic analyses (e.g. also performing functional or assembly analyses, such as Morais et al. 2022; Boulund et al. 2023) of which each step has fewer options of tools and may execute functionality unwanted by the end user.

Here we present nf-core/taxprofiler (https://nf-co.re/taxprofiler), a pipeline designed to allow users to efficiently and simultaneously taxonomically classify or profile short- and long-read sequencing data. At the time of writing it supports 11 classifiers and an arbitrary number of databases per classifier in a single pipeline run. nf-core/taxprofiler utilises Nextflow (Di Tommaso et al. 2017) to ensure efficiency, portability, and scalability, and has been developed within the nf-core initiative of Nextflow pipelines (Ewels et al. 2020) to ensure high quality coding practises and user accessibility. It includes detailed documentation and a graphical-user-interface GUI) execution interface in addition to a standard command-line-interface (CLI).

## 3 Description

nf-core/taxprofiler aims to facilitate three main steps of a typical whole-genome, metagenomic sequencing analysis workflow (Chiu and Miller 2019,Figure 1). A longer description of the available functionality and motivations can be seen in the Supplementary Information.

**Figure 1:**
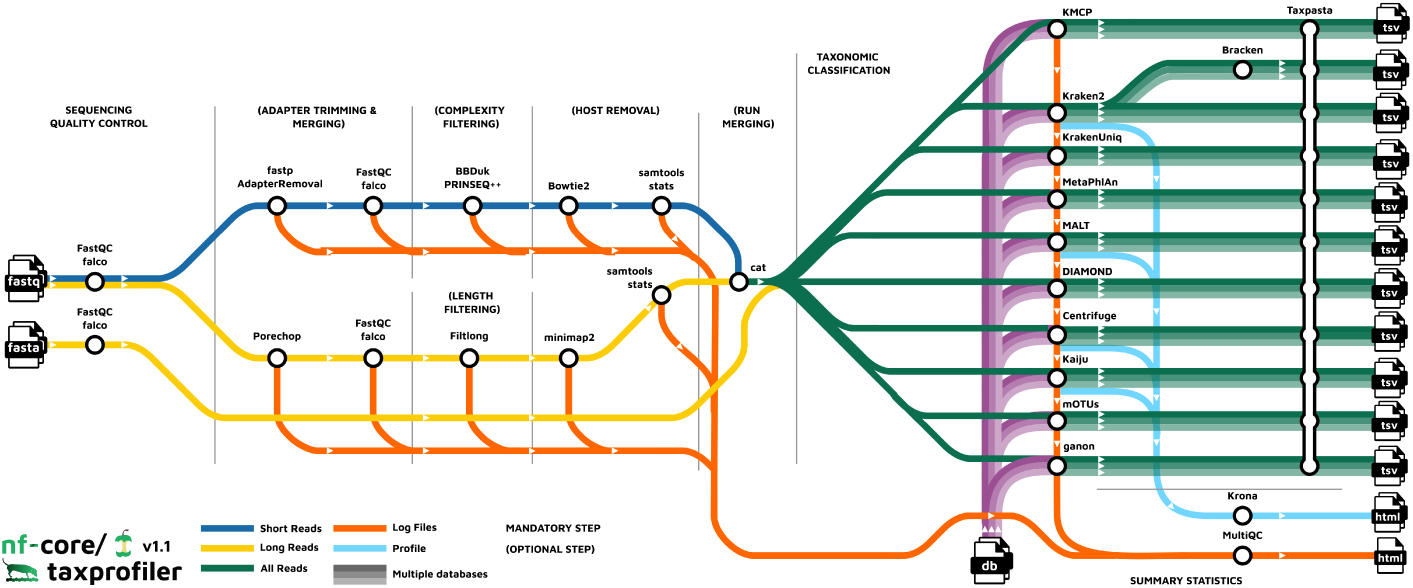
Visual overview of the nf-core/taxprofiler workflow. nf-core/taxprofiler can take in FASTQ (short or long reads) or FASTA files (long reads), that will optionally go through sequencing quality control (e.g. with FastQC), read preprocessing e.g. removal of adapters), complexity filtering, host removal, and run merging before performing taxonomic classification and/or profiling with a user-selected range of tools and databases. Output from all classifiers and profilers are standardised into a common taxon table format, and when supported visualisations of the profiles are generated.

In brief, nf-core/taxprofiler can accept short-(e.g. Illumina) and/or long-read e.g. Nanopore) FASTQ or FASTA files. These are supplied to the pipeline in the form of a TSV file that includes basic sample and sequencing library metadata. The pipeline can then be executed either via a standard Nextflow command-line-interface execution or graphical-user-interface through either the open-source and free nf-core launch page (https://nf-co.re/launch) or the commercial (with free-tier) Nextflow tower (https://tower.nf) solution. Examples of the command-line execution and nf-core launch GUI can be seen in the Supplementary Information.

The pipeline can perform a range of metagenomics appropriate read preprocessing steps, such adapter removal, read merging, low-sequence complexity filtering, host-or contamination removal, and/or per-sample run merging. All of these steps are optional, and are aimed at removing possible sequencing artefacts that may result in false positive taxonomic classification hits or improve classification efficiency. Most of these steps also provide options of different tools to account for user preference.

After pre-processing, nf-core/taxprofiler can perform simultaneous profiling of pre-processed reads with up to as many as 11 different taxonomic classifiers or profilers Table 1). Additionally on top of this, also simultaneously for each of the classifiers, an arbitrary number of databases as supplied by the user. As of version 1.1.0, the following classifiers and profilers are available: Kraken2 (Wood, Lu, and Langmead 2019), Bracken (Lu et al. 2017), KrakenUniq (F. P. Breitwieser, Baker, and Salzberg 2018), Centrifuge (Kim et al. 2016), MALT (Vågene et al. 2018), DIAMOND (Buchfink, Reuter, and Drost 2021), Kaiju (Menzel, Ng, and Krogh 2016), MetaPhlAn (Blanco-Míguez et al. 2023), mOTUs (Ruscheweyh et al. 2022), ganon (Piro et al. 2020), and KMCP (Shen et al. 2023). Databases are also supplied via a input TSV file, which also allows per-database custom classification parameters - meaning a given database can be supplied multiple times each with different parameters or multiple different databases per profiler. All classifiers with secondary steps to generate or convert to additional output file formats are also included.

**Table 1:**
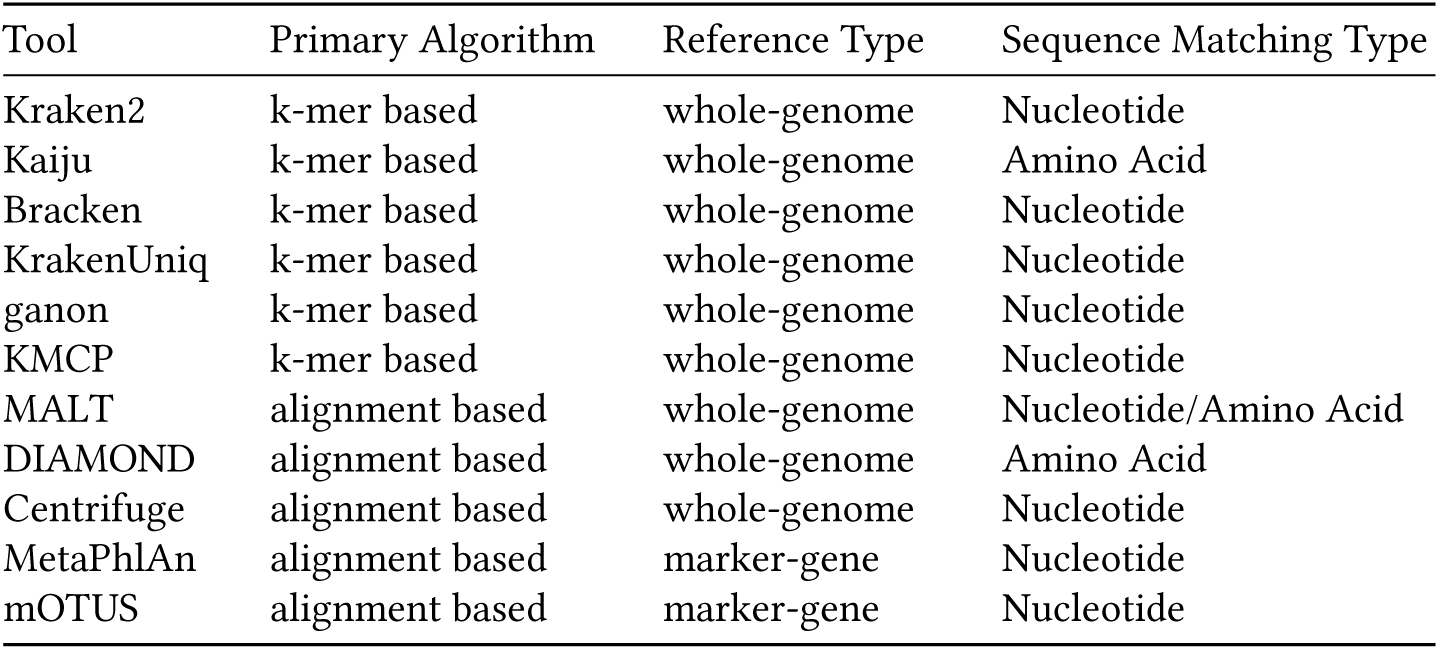
List of nf-core/taxprofiler supported taxonomic/classifiers profilers as of version 1.1 and their approximate method and supported input database types. Primary algorithm refers to the algorithm type used for sequencing matching. Reference type refers to the typical sequence type used in database construction of the tool. Sequencing matching type refers to which ‘molecular alphabet’ is primarily used for matching between a query (read) and a reference (genome/gene).

| Tool | Primary Algorithm | Reference Type | Sequence Matching Type |
| --- | --- | --- | --- |
| Kraken2 | k-mer based | whole-genome | Nucleotide |
| Kaiju | k-mer based | whole-genome | Amino Acid |
| Bracken | k-mer based | whole-genome | Nucleotide |
| KrakenUniq | k-mer based | whole-genome | Nucleotide |
| ganon | k-mer based | whole-genome | Nucleotide |
| KMCP | k-mer based | whole-genome | Nucleotide |
| MALT | alignment based | whole-genome | Nucleotide/Amino Acid |
| DIAMOND | alignment based | whole-genome | Amino Acid |
| Centrifuge | alignment based | whole-genome | Nucleotide |
| MetaPhlAn | alignment based | marker-gene | Nucleotide |
| mOTUS | alignment based | marker-gene | Nucleotide |

**Table 2:**
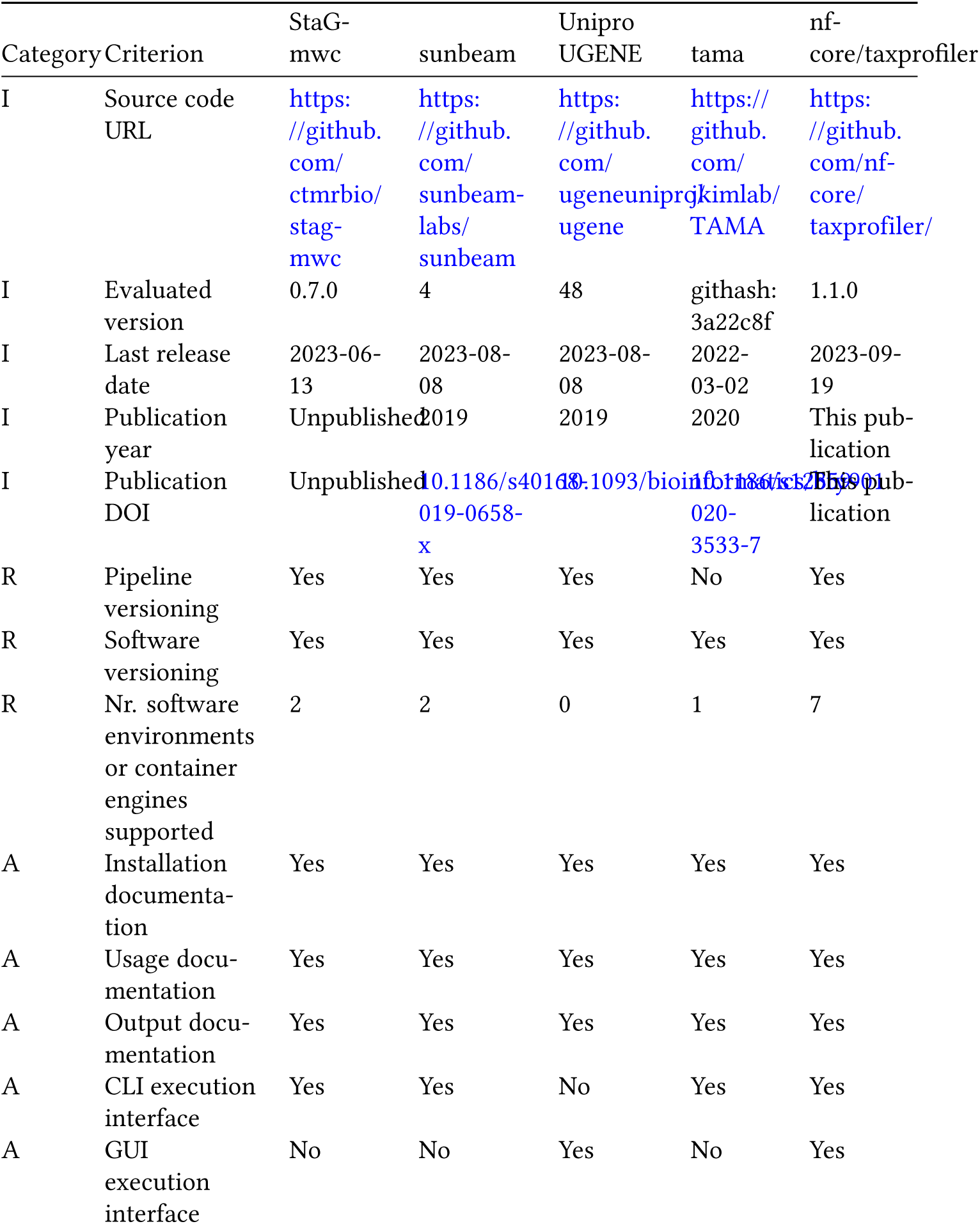
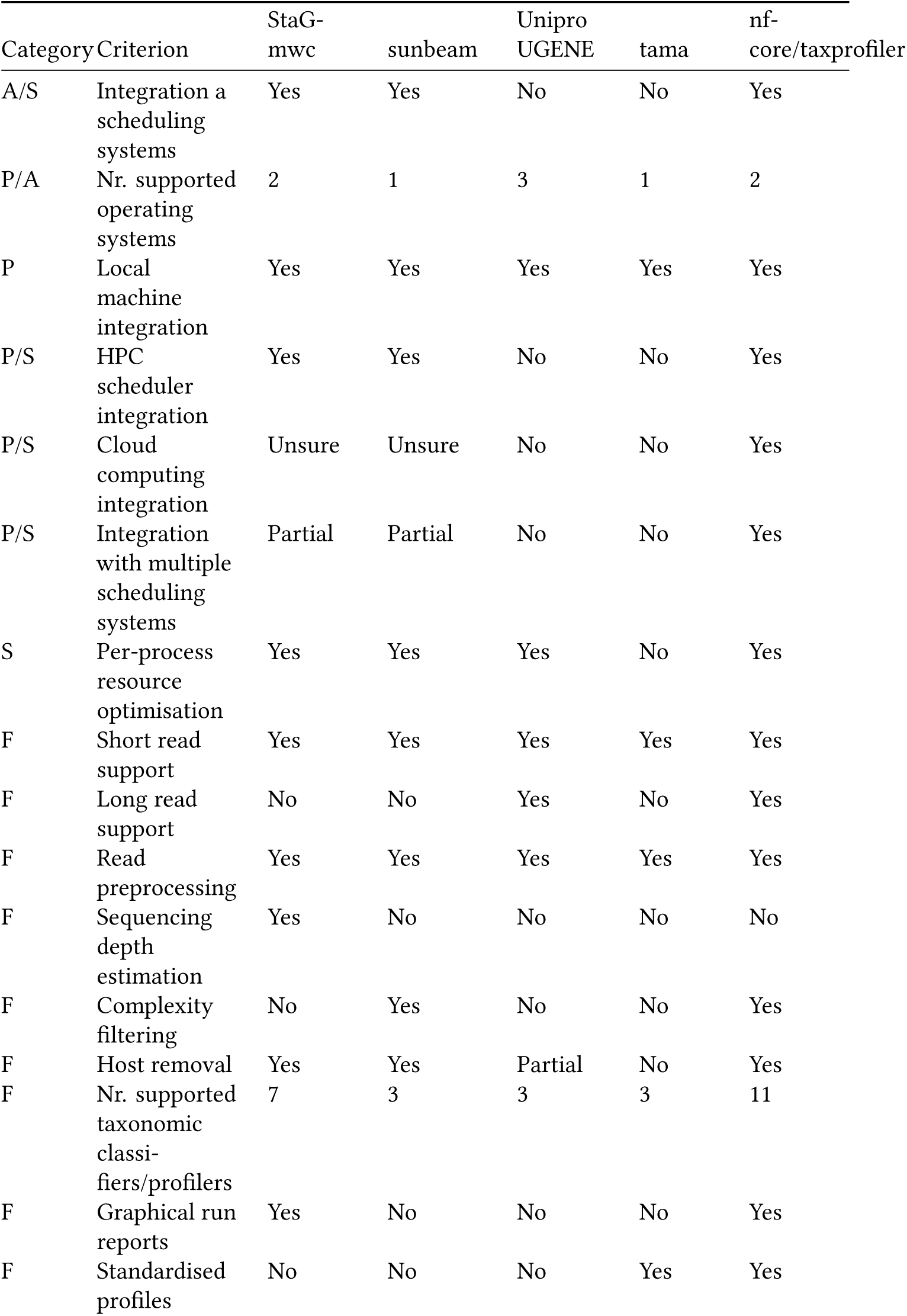
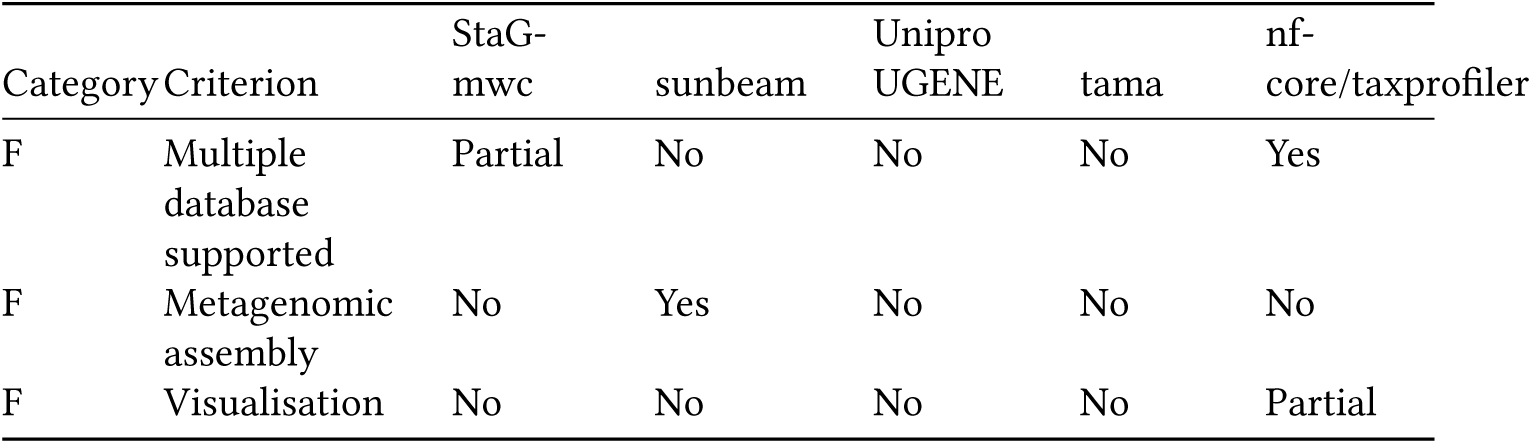
Comparison of functionality with four recent taxonomic pipelines with similar functionality. A more detailed textual comparison can be found in the Supplementary Information. Category keys are as follows: I - Information, R - Reproducibility, A - Accessibility, P - Portability, S - Scalability, F - Functionality.

Post-processing of taxonomic profiles include standardisation and aggregation of profiles, i.e. merging of multiple profiles into a single multi-sample table for easier comparison between profilers, with the tool TAXPASTA (Beber et al. 2023), and visualisation of profiles with Krona (Ondov, Bergman, and Phillippy 2011) where supported.

All relevant preprocessing statistics are displayed in an interactive and dynamic MultiQC report (Ewels et al. 2020).

nf-core/taxprofiler comes with extensive documentation for general usage, short- and long-parameter help texts, and output file descriptions. To ensure maximum accessibility, these are available in pipeline results as markdown files (https://github.com/nf-core/taxprofiler), on the nf-core website (https://nf-co.re/taxprofiler) and for the parameter help texts on the command line via standard --help. The output documentation also aims to guide users as the most suitable files for different types of downstream analysis

## 4 Discussion

A range of pipelines already exists for taxonomic profiling, however, each have their own particular purpose and capabilities. We compared the functionality of nf-core/taxprofiler against four other recently published or released pipelines, selected based on their similarity of purpose to nf-core/taxprofiler. The selection criteria and a more detailed comparison between the five pipelines can be seen in the Supplementary Information. Overall, while there was a general similarity across all pipelines, nf-core/taxprofiler showed the largest number of options for pipeline execution accessibility, and user choice. This is facilitated through the use of an established workflow manager (with Nextflow supporting 7 software environment/container systems), support for both CLI and GUI execution, and by the number of supported classifiers. Furthermore, it is unique in that is the only pipeline to support supplying multiple database for all of the tools in a single pipeline run.

Another important advantage of nf-core/taxprofiler is that it is being developed within the nf-core community (https://nf-co.re), that provides strong long-term support for the continued community-based development and maintenance of its pipelines. In this framework, we will continue to add additional preprocessing, metagenomic classification, and profiling tools as they become established and as requested by the metagenomics community. For example, we feel that the inclusion of steps such as sequencing saturation estimation as already being performed by a similar pipeline StaG-mwc (https://github.com/ctmrbio/stag-mwc) would be beneficial to the nf-core/taxprofiler workflow (possibly with dedicated tools such as Nonpareil, Rodriguez-R et al. 2018), and/or more performant complexity filtering tools such as Komplexity as offered by the sunbeam metagenomics pipeline (Clarke et al. 2019). Additional tools that could be added for short-read classification could include sourmash (Titus Brown and Irber 2016) that provides scalable sequence to sequence comparison or other marker gene reference tools such as tools such as METAXA2 (Bengtsson-Palme et al. 2015) that use shotgun sequencing reads to recover 16S sequences from metagenomic samples. Adding additional classifiers also applies to extend support to other sequencing platforms; nf-core/taxprofiler already supports Nanopore long-read data, however the use of long-read PacBio data for metagenomic data is growing in interest (Portik, Brown, and Pierce-Ward 2022). We are therefore considering adding dedicated preprocessing steps for this type of sequencing data.

A remaining major challenge for metagenomics researchers (and not supported in the same workflow by any of the compared pipelines above) is the construction of databases for each profiling tool. Given there still are no curated, high-quality ‘gold standard’ databases in metagenomics, and while nf-core/taxprofiler allows the profiling against multiple databases and settings in parallel, currently the pipeline still requires users to construct these manually and to supply to the pipeline. While we feel this is currently a reasonable investment as such databases are typically repeatedly re-used, we are exploring the possibility to add an additional complementary workflow in the pipeline to allow automated database construction of all classification tools, given a set of FASTA reference files.

Finally, once an overall taxonomic profile is generated, researchers often wish to validate hits through more sensitive and accurate methods such as with read-mapping alignment. While read alignment is supported by other pipelines such as StaG-mwc, this happens in-parallel to the taxonomic profiling and requires prior expectation of which reference genomes to map against. Instead, nf-core/taxprofiler could be easily extended to have a validation step similar to the approach of the ancient DNA metagenomic pipeline aMeta (Pochon et al. 2022). Utilising Nextflow’s execution parallelism, the input sequences could be aligned back to the reference genomes of only those species with hits resulting from the taxonomic classification, but with dedicated accurate short- or long-read aligners. In addition to the more precise classification, post-classification read-alignment could also be particularly useful for researchers in palaeogenomics who wish to use tools other than KrakenUniq for initial classification as in aMeta), where alignment information can be used to authenticate ancient DNA within their samples, but also in clinical metagenomics to identify potential pathogens at much finer resolution (e.g. down to strain level).

Another motivation for developing nf-core/taxprofiler, despite the large number of existing metagenomics pipelines, is that by establishing a taxonomic profiling pipeline within the nf-core ecosystem, it is possible to begin building both standalone but also an integrated suite of powerful interconnected pipelines for the major stages of metagenomic workflows. Existing microbial- and metagenomics-related pipelines within the nf-core initiative include nf-core/ampliseq (Straub et al. 2020), nf-core/mag Krakau et al. 2022), and nf-core/funcscan (https://nf-co.re/funcscan). We expect over time the ability to link inputs and outputs of each workflow to develop comprehensive metagenomic analyses, while still maintaining powerful standalone pipelines, providing maximal user choice but with familiar interfaces.

## 5 Conclusion

nf-core/taxprofiler is an accessible, efficient, and scalable pipeline for metagenomic taxonomic classification and profiling that can be executed on anywhere from laptops to the cloud. To our knowledge, the pipeline offers the largest number of taxonomic profilers across similar pipelines, providing flexibility for users not just on choice of profiling tool but also with databases and database settings within a single run. With the development within the open and welcoming nf-core community and with best-practise development infrastructure, we look forward to further contributions and involvement of the wider metagenomics community, and also we hope that through detailed documentation and a range of execution options, nf-core/taxprofiler will make reproducible and high-throughput metagenomics more accessible for a wide range of disciplines.

## 6 Code Availability

nf-core/taxprofiler source code is available on GitHub at https://github.com/nf-core/ taxprofiler, and each release is archived on Zenodo (latest version DOI: 10.5281/zenodo.7728364)

The version of the pipeline described in this paper is version 1.1.0 (release specific Zenodo archive DOI: 10.5281/zenodo.8358147)

## 7 Acknowledgments

We thank Prof. Christina Warinner and the Microbiome Sciences group MPI-EVA for original discussions that lead to the pipeline. We are also grateful for the nf-core community for the original and ongoing support in the development in the pipeline, in particular for the contributions by Lauri Mesilaakso, Jianhong Ou, and Rafał Stępień.

## 10 Supplementary Information

### 10.1 Implementation

#### 10.1.1 Input and Execution

The pipeline can be executed via typical Nextflow commands (Code Block 1), or using the standard nf-core ‘launch’ GUI (Figure 2), making the pipeline accessible for both computationally experienced as well as less experienced researchers. In addition to the general usage and parameter documentation of the pipeline (https://nf-co.re/taxprofiler). The GUI offers immediate assistance and guidance to users on what each parameter does, both in short- and long-form, with long-form parameter descriptions additionally describing which tool-specific parameters are being modified for each pipeline parameter (https://nf-co.re/launch/?pipeline=taxprofiler). The GUI also includes controlled user input by providing strict drop-down lists and input validation prior execution of the pipeline (Figure 2) to reduce the risk of typos and other mistakes, which is in contrast to the command-line interface that only includes validation at pipeline run-time.

**Figure 2:**
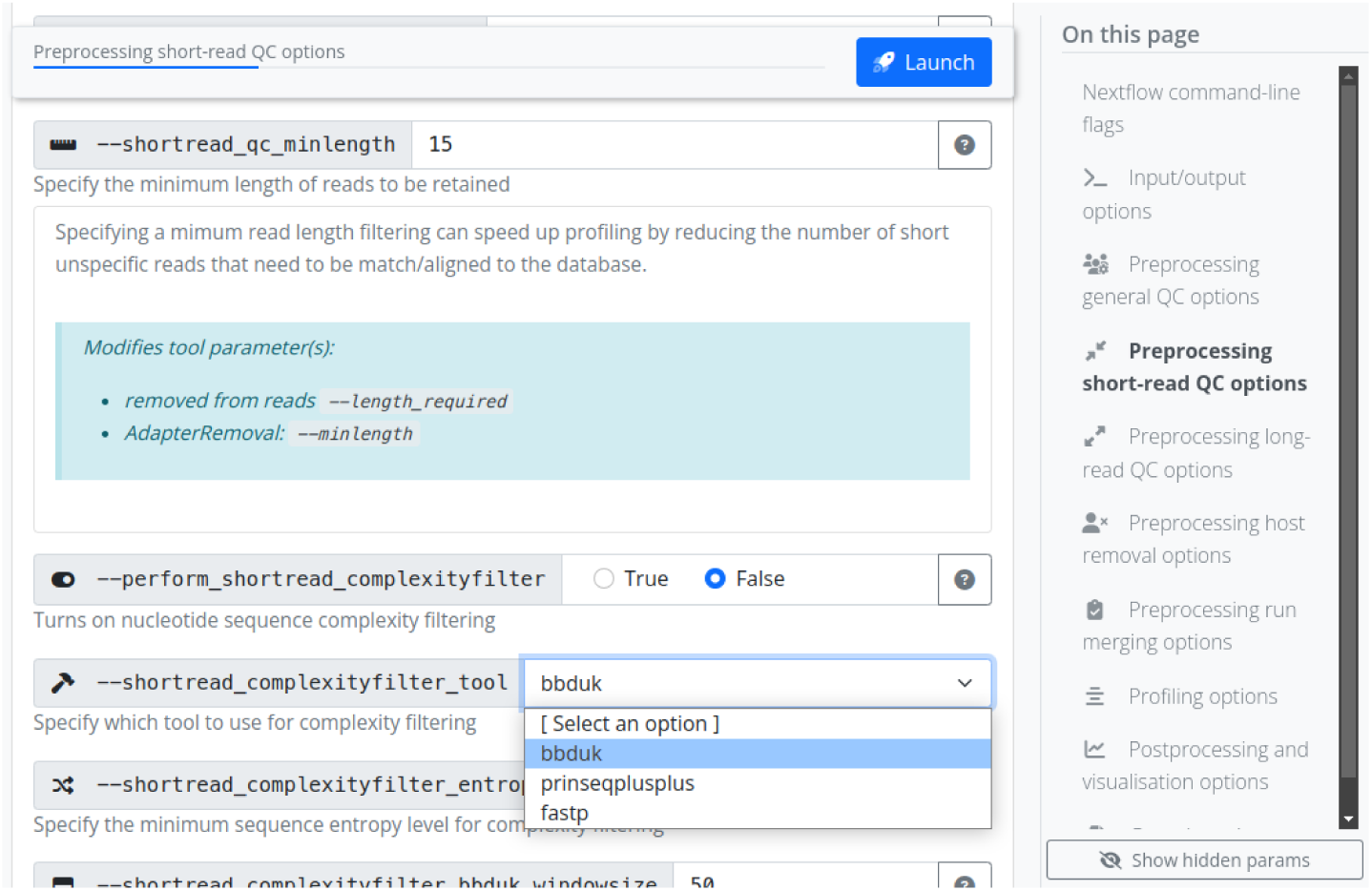
Screenshot of the nf-core pipeline launch graphical user interface with nf-core/taxprofiler options displayed. The web browser-based interface provides guidance for how to configure each pipeline parameter by providing both short and long help descriptions to help guide users in which contexts to configure each parameter. Additional elements such as radio buttons, drop down menus, and background regular expressions check for validity of input. When pressing launch, a prepared configuration file and command is provided that can be copied and pasted by the user into the terminal with the corresponding pipeline parameter (e.g. --run_kraken2). Per-classifier flags are also available for the optional saving of additional non-profile output files. Alternatively to command line flags, parameters can be specified via pre-configured YAML format files, with which (provided no hardcoded paths are included) can be re-used across pipeline runs.

**Listing 1.**
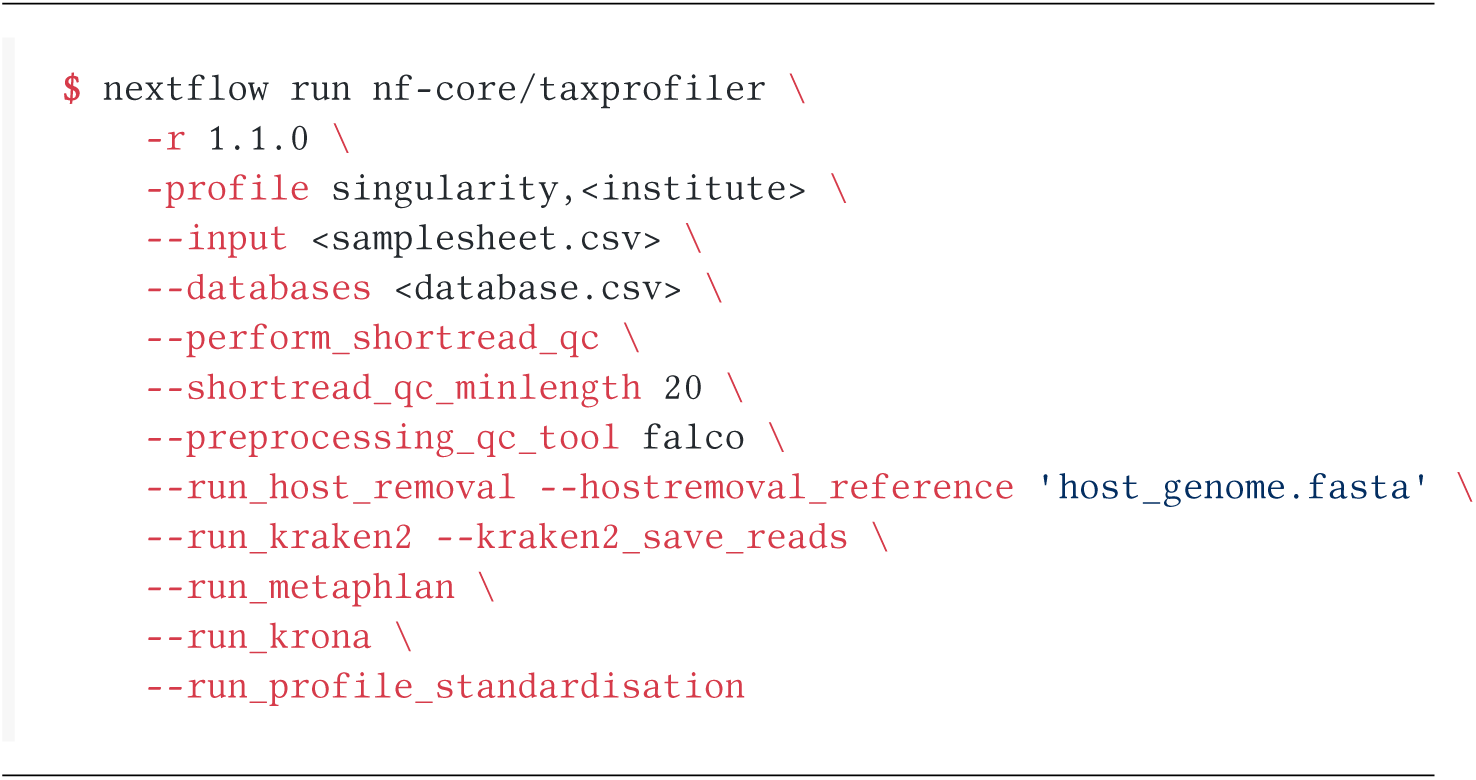
Example nf-core/taxprofiler command for running short-read quality control, removal of host DNA and executing the k-mer based Kraken2 and marker gene alignment MetaPhlAn tools.

An example nf-core command line execution of the pipeline can be seen in Code Block 1, where two input files are supplied: one file specifying paths of FASTQ files of metagenomic samples and necessary metadata for preprocessing (such as sample ID and sequencing platform), and the second file specifying paths to the user-defined databases with per-database classification parameters. Various parameters are available to select different preprocessing steps, and provide additional configuration such as tool selection and value options. Note that even if a user supplies a given database in the database input sheet, the corresponding profiling tool must still be activated

All nf-core pipelines are strictly versioned (specified with the Nextflow -r flag), and to ensure reproducibility, each version of the pipeline has a fixed set of software used for each step of the pipeline. The fixed set of software are controlled through the use of the conda package manager or containers (Docker, or Apptainer - previously known as Singularity, etc) from the stable Bioconda (Grüning et al. 2018) or BioContainers Veiga Leprevost et al. 2017) repositories. This, coupled with the intrinsic Nextflow ability to execute on most infrastructure whether that is a local laptop (resource requirements permitting), traditional HPC, as well across common cloud providers also makes nf-core/taxprofiler a very portable pipeline that can be used in many contexts.

#### 10.1.2 Preprocessing

Preprocessing steps in nf-core/taxprofiler are aimed at removing laboratory and sequencing artefacts that may influence taxonomic profiling, either for computing resource consumption or and/or false-positive or false-negative classification reasons. First sequencing quality control with FastQC (Andrews 2010) or Falco (Sena Brandine and Smith 2021) is carried out. Falco was included for reduced memory requirements, in particular for long read sequencing data. Artificial library adapter sequences added during sequencing reduce sequencing matching accuracy by reducing sequence specificity, and in some cases, may result in false-positive hits due to adapter sequence contamination in reference genomes (Schäffer et al. 2018; F. P. Breitwieser, Baker, and Salzberg 2018) ^1^. Additionally, paired-end merging may provide longer sequences that will allow for more specific classification when paired-end alignment is not supported by a given classifier. For these tasks nf-core/taxprofiler can apply either fastp Chen et al. 2018) or AdapterRemoval2 (Schubert, Lindgreen, and Orlando 2016) for short reads, and currently Porechop (Wick et al. 2017) for Oxford Nanopore long-read data. For both short and long reads, FastQC or Falco is run again to allow assessment on the performance of the adapter removal and/or pair-merging step.

Low complexity sequences, e.g. sequences containing long stretches of mono- or di-nucleotide repeats provide little specific genetic information that contribute to taxonomic identification, as they can align to many different reference genomes Schmieder and Edwards 2011; Clarke et al. 2019). Including such reads during taxonomic profiling can increase run-time and memory usage for little gain, as during lowest-common-ancestor (LCA) classification steps they will be assigned to high-level taxonomic ranks (e.g. Kingdom). nf-core/taxprofiler performs removal of these reads through complexity filtering algorithms as provided by fastp, BBDuk Bushnell 2022), or PRINSEQ++ (Cantu, Sadural, and Edwards 2019). Long read sequences often do not have such reads, as lengths are sufficient enough to capture greater sequence diversity - but it is sometimes desirable to only classify reads longer than a certain length - as these provide more precise taxonomic information (Dilthey et al. 2019; Portik, Brown, and Pierce-Ward 2022). Therefore, nf-core/taxprofiler can remove reads shorter than a user-defined length using Filtlong.

Removing host DNA is another common preprocessing step in metagenomic studies. This can help speed up run-time, particularly in microbiome studies, where detection of microbes are of interest. Furthermore, host-contamination of reference genomes in public databases is common (Longo, O’Neill, and O’Neill 2011; Kryukov and Imanishi 2016; Florian P. Breitwieser et al. 2019). Therefore, the removal of such sequences can help decrease the risk of false positive taxonomic assignment. To remove multiple hosts or other sequences, all reference genomes can be combined into a single FASTA reference file. Short read host removal can be carried out with Bowtie2 (Langmead and Salzberg 2012; Langmead et al. 2019) and minimap2 (Li 2018) for long reads, both in combination with SAMtools (Li et al. 2009; Danecek et al. 2021), where reads are aligned against the reference genome and the off-target (unaligned) reads are then converted back to FASTQ format for classification.

Finally, nf-core/taxprofiler can optionally perform ‘run merging’ where multiple FASTQ files from the same sample but have been sequenced over multiple lanes are concatenated together to generate one profile per sample or library. The final set of reads used for profiling can be optionally saved for downstream re-use. Throughout all steps, relevant statistics and log files are generated and used both for the final pipeline run report as well as saved into the results directory of the pipeline run for further inspection where necessary.

#### 10.1.3 Profiling

There are many types of metagenomic profiling techniques, from profiling against whole-genome references with alignment or k-mer based approaches, to methods involving alignment to species-specific marker-gene families (Quince et al. 2017; Ye et al. 2019). nf-core/taxprofiler aims to support and include all established classification or profiling tools as requested by the community.

The choice of tools used in a pipeline run is up to the user, with a tool being executed when both the corresponding database and --run_<tool> flag is provided. Specific classification settings for each tool and database are specified in the database CSV input sheet. Some tools also have pipeline level command-line flags for controlling certain aspects of output files.

The following classifiers and profilers are supported in version 1.1.0 of nf-core/taxprofiler: Kraken2 (Wood, Lu, and Langmead 2019), Bracken (Lu et al. 2017), KrakenUniq (F. P. Breitwieser, Baker, and Salzberg 2018), Centrifuge (Kim et al. 2016), MALT (Vågene et al. 2018), DIAMOND (Buchfink, Reuter, and Drost 2021), Kaiju (Menzel, Ng, and Krogh 2016), MetaPhlAn (Blanco-Míguez et al. 2023), mOTUs Ruscheweyh et al. 2022), ganon (Piro et al. 2020), KMCP (Shen et al. 2023).

By default, nf-core/taxprofiler produces the default per-sample taxonomic classification profile output from a tool or a tool’s report generation tool. The output is normally in the form of counts per reference sequencing, with additional statistics about the hits of a particular organism (estimated sequence abundance, taxonomic level etc.). Users can also optionally request output of per-read classification output and output such as classified and unclassified reads in FASTQ format, where supported.

The pipeline provides high efficiency, particularly during the metagenomic classification stage, through the inherent parallelisation provided by Nextflow. While metagenomic classification is comparatively computationally intensive (in terms of memory and execution time; due to a combination of sequencing depth and number of reference genomes), Nextflow automatically optimises the execution order of all the steps in pipeline, maximising the number parallel running of multiple profilers and/or databases at any given time point, as far as the available computational resources allow. For local machines such as laptops or desktops, Nextflow will automatically detect all available computational resources, but this is customisable using Nextflow configuration files. For HPC and cloud infrastructure, users typically have to define the computational infrastructural environment the pipeline is being executed on (CPU or memory limitations, queues, instance types, etc.). To facilitate the pipeline computational configuration, nf-core/taxprofiler supports use of more than 90 pre-defined centralised generic and pipeline-specific institutional Nextflow configurations as provided by nf-core/configs (https://nf-co.re/configs). However, of course users are still welcome to supply their own custom configuration files as with any typical Nextflow run, further refining computational limitations or execution specifications.

### 10.1.4 Post-profiling

In metagenomic studies, it is common practise to compare the profiles among many samples, and the results of multiple profiles are normally stored in ‘taxon tables’, i.e, counts per reference taxon (rows), for each sample (columns). When available, nf-core/taxprofiler supports the option to produce the ‘native’ taxon table of each classification tool when multiple samples are run.

One of the challenges that researchers face when comparing multiple taxonomic classifiers or profilers is the heterogenous output formats that are produced, that often require custom parsing and merging scripts for each tool to standardise. To facilitate more user-friendly cross-comparisons between tools, nf-core/taxprofiler utilises the TAXPASTA tool (Beber et al. 2023) to generate standardised profiles and generate multi-sample tables.

Summary statistics for the entire pipeline are visualised and displayed in a customisable MultiQC report (Ewels et al. 2020). When supported, quality control of data and pipeline runs are shown for manual verification. Krona plots (Ondov, Bergman, and Phillippy 2011) can also optionally be generated for supported tools to help provide further visualisation of taxonomic profiles.

#### 10.1.5 Output

To summarise, the main default output from nf-core/taxprofiler are both classifier native’ and standardised single- and multi-sample taxonomic profiles with counts per-taxon and an interactive MultiQC run report with all run statistics, in addition to the raw log files themselves where available.

The MultiQC run report displays statistics and summary visualisations for all steps of the pipeline where possible, lists of versions for all tools of each step of the pipeline. It also provides a dynamically-constructed text for the recommended ‘methods’ for reporting how the pipeline was executed (including relevant citations) that users can use in their own publications.

Optional outputs can include other types of profiles (e.g. per read classification) and in other formats as produced by the tools themselves, as well as raw reads from pre-processing steps and output visualisations from Krona. Nextflow resource usage and trace reports are also by default produced for users to check pipeline performance.

### 10.2 Comparison with other solutions

nf-core/taxprofiler has been specifically developed for the analysis of whole-genome, *metagenomic* sequencing data. While other types of taxonomic profiling data such as 16S amplicon sequencing are well established fields with a range of popular high-quality and best-practise tools pipelines (e.g. Blanco-Míguez et al. 2023; Schloss et al. 2009) and databases (DeSantis et al. 2006; Yilmaz et al. 2014), ‘gold standard’ tools and databases for metagenomics remain much less established. Thus, the need for highly-multiplexed classification is more desirable for the newer metagenomics methods.

We searched Google Scholar for open-source pipelines published or released in the last 5 years (at the time of writing, since 2018) that were designed primarily for metagenomic classification screening, that supported at least 2 classifiers, had at least one preprocessing step and were not specifically targeted at read classification of specific domains of taxa (e.g. viruses or bacteriophages only). We also included an additional open-source but unpublished pipeline at the recommendations of the authors of the pipeline due to the functional overlap to nf-core/taxprofiler. We then evaluated the pipelines based on their publications and documentation for typical metagenomic profiling workflow steps. We used a range of criteria related to expectations of modern bioinformatic workflows that can be summarised in the following four categories: reproducibility, accessibility, scalability, and portability (Wratten, Wilm, and Göke 2021). After searching, we selected the following pipelines for comparison with nf-core/taxprofiler that matched the specific criteria described above: sunbeam (v4, Clarke et al. 2019), Unipro UGENE (v48, Rose et al. 2019), TAMA (githash: 3a22c8f, Sim et al. 2020), and StaG-mwc (0.7.0, Boulund et al. 2023).

In terms of accessibility, all pipelines have documentation describing the installation steps, usage instructions, and output files. However, there are varying levels of detail and comprehensiveness. In particular, StaG-mwc and nf-core/taxprofiler have the most detailed descriptions of all possible output files for every supported module, whereas Unipro UGENE and sunbeam have very minimal to possibly unfinished output documentation. For execution options, most of the pipelines provide CLI execution, except for Unipro UGENE which offers only GUI-based pipeline set-up (despite a command-line execution of the GUI generated configuration). In particular, nf-core/taxprofiler is the only pipeline providing both CLI and GUI interfaces for pipeline run execution.

Criteria covering portability also overlap with accessibility, as it implies options for and ease of different users running on different types of computing infrastructure, whether that is on their own laptop, on an HPC cluster, or in the cloud. Unipro UGENE is the only pipeline that explicitly satates support for execution on all three major operating systems (Linux, OSX, Windows), whereas StaG-mwc and nf-core/taxprofiler can be run on unix operating systems (albiet possibly on Windows via Windows Sub-system for Linux (WSL)), and sunbeam and TAMA are only being supported on Linux.

While all pipelines support ‘local’ machine execution (e.g. personal laptops or desktops), a large portion of academic users execute computationally intensive bioinformatic tasks on HPC clusters. In these contexts, pipeline task submissions are normally managed by job schedulers, thus integration with schedulers is an important criterion for running large multi-step and parallelised pipelines. The three pipelines leveraging workflow managers (Snakemake and Nextflow) support integration with schedulers StaG-mwc, sunbeam, and nf-core/taxprofiler) with nf-core/taxprofiler supporting the most by far (>10 scheduling systems) as natively offered by Nextflow. This allows the greatest possible choice for users in terms of which HPC infrastructure they can execute their pipeline on. As an extension of this, only nf-core/taxprofiler has explicit support for cloud computing (e.g. AWS, GCP, or Microsoft Azure) as provided by Nextflow, again maximising user choice and portability when it comes to running the pipeline.

In terms of scalability, the aforementioned integration with schedulers and cloud computing support implicitly maximises efficiency and parallellisation of pipeline runs, providing good scalability for varying numbers of input files and steps in the pipeline. Again, the three workflow manager based pipelines provide scalability, whereas there is no mention neither Unipro UGENE nor TAMA in reference to parallel task execution. Furthemore, all pipelines except TAMA, allowed per-process customisation of computational resources, something critical for maximising efficient scalability to ensure only the necessary resources for a given step of a pipeline are requested.

In terms of reproducibility, all five pipelines are good at ensuring reproducibility in terms of pipeline and software versioning (allowing re-execution of pipeline runs using the same software), with only TAMA not having stable versioned releases. However, installing software manually across different infrastructures can result in variability in the execution of each software ^2^ (Di Tommaso et al. 2017).The current most popular solution to the problem of inconsistent software environments is to use container engines such as Docker or Apptainer to run container images which are isolated, deterministic computing environments which can be executed by any system providing a container runtime. Only Unipro UGENE does not document the use of a container system, with nf-core/taxprofiler offering the biggest choice for users, again, courtesy of Nextflow with 6 different engine systems at the time of writing.

Finally, we compared metagenomics related functionality between the pipelines. All pipelines support short-read FASTQ input, but only nf-core/taxprofiler explicitly reports long-read support, while the documentation in Unipro UGENE states that assembled contigs are possible input to some of the profilers. All pipelines support read pre-processing (adapter clipping, and merging). In terms of tools used for preprocessing, Trimmomatic (Bolger, Lohse, and Usadel 2014) is popular across the other pipelines but is not supported in nf-core/taxprofiler. Only sunbeam and nf-core/taxprofiler support complexity filtering to remove low sequence diversity reads. In fact within sunbeam, the authors developed their own dedicated, performant complexity filtering tool Komplexity (Clarke et al. 2019). Most pipelines support some form of host removal (only TAMA did not support this), and it is likely possible with Unipro UGENE although not directly described). In all cases, host removal consists of mapping processed reads with an aligner and using the off-target reads for downstream profiling as implemented in nf-core/taxprofiler), however StaG-mwc has an additional separate metagenomic host removal step with Kraken2. nf-core/taxprofiler supports by far the largest number of taxonomic classifers and profilers at 11 as of v1.1.0 - providing the greatest choice to users - with StaG-mwc offering 7, and the remaining pipelines only 3. Only nf-core/taxprofiler and partly StaG-mwc explicitly support running each profiler with multiple databases. nf-core/taxprofiler is the only pipeline that supports running an arbitrary number of different metagenomic profiler databases each with their own settings. This makes it a useful for tool parameter comparison, testing different databases, or reducing the size of each database (e.g. per domain) to make it more flexibility for running on smaller computational infrastructure. StaG-mwc allows multiple references for their short-read alignment steps rather than the metagenomic profilers. For output, nf-core/taxprofiler, StaG-mwc, and sunbeam via an extension) support a singular run report for summarising all preprocessing step. Only nf-core/taxprofiler and TAMA produce standardised output for all taxonomic profilers, the former with the dedicated standalone tool TAXPASTA (Beber et al. 2023). However Unipro UGENE additionally offers a ‘consensus’ profile using WEVOTE (Metwally et al. 2016).

To summarise, many of the pipelines reviewed here offer similar functionality, with particularly StaG-mwc having a strong overlap with nf-core/taxprofiler. Thus, users in most cases will be able to select the pipeline depending on which framework they feel most comfortable with. However the advantages of nf-core/taxprofiler mainly come from the offering of the greatest choice of tools, as well the particular benefits provided by Nextflow. It provides the greatest number of computational infrastructure types the pipeline can be executed on, and container systems can be used to ensure reproducibility, as well the support of the nf-core community due to the centralised pool of ‘plug-and-play’ modules to make it easier to update the pipeline over time to add new tools classifiers.

The functionality offered by other pipelines not currently supported by nf-core/taxprofiler include sequencing saturation estimation (StaG-mwc), taxonomy-free composition comparison (StaG-mwc), functional profiling (StaG-mwc), *de novo* assembly (sunbeam), and reference mapping (StaG-mwc, sunbeam). We do not plan to support *de novo* assembly or functional profiling in nf-core/taxprofiler as we feel these are already better served by other existing dedicated pipelines within the nf-core ecosystem: nf-core/mag for *de novo* assembly, (Krakau et al. 2022) and nf-core/funcscan for functional profiling (https://nf-co.re/funcscan), as well as elsewhere e.g. MetaWRAP (Uritskiy, DiRuggiero, and Taylor 2018).

## Footnotes

1 For an ‘infamous’ case of adapter sequences in a published eukaryotic genome, see the following blog posts Graham Etherington: https://web.archive.org/web/20201219022000/http://grahametherington.blogspot.com/2014/09/why-you-should-qc-your-reads-and-your.html?m=1why-you-should-qc-your-reads-and-your.html Sixing Huang: https://web.archive.org/web/20220904205331/https://dgg32.medium.com/carp-n-the-soil-1168818d2191 (Accessed 2023-08-25)

2 As demonstrated in this blogpost from Paweł Przytuła: https://web.archive.org/web/20230320223436/ https://appsilon.com/reproducible-research-when-your-results-cant-be-reproduced/ (Accessed 2023-08-25)

